# The Evolution of Immune Sensitivity under Immunopathological and Autoimmune Costs

**DOI:** 10.1101/2020.10.12.335620

**Authors:** Lafi Aldakak, Frank Rühli, Matthias Galipaud, Nicole Bender

## Abstract

Hosts with high immune sensitivity benefit from rapid recovery but suffer multiple costs thereof. We distinguish between immunopathological costs due to collateral damage and autoimmune costs due to false positives against self. Selection on sensitivity follows different trajectories depending on the cost nature and the overlap degree between host- and parasitic molecular signatures. Increased parasite virulence selects for higher immune sensitivity under immunopathological costs but low sensitivity when the costs are autoimmune, contradicting previous theoretical results. Longer lifespan of the host selects for low sensitivity under immunopathology to avoid accumulated tissue damage. Under autoimmune costs, hosts with a shorter lifespan cannot afford to shorten it further due to autoimmunity and evolve lower immune sensitivity. Longer lifespan selects for high or low sensitivity depending on the presence of immune memory. These results extend our understanding of selection on immune sensitivity and help explain phenomena like the cytokine shock and chronic infections.

## 1. Introduction

Parasites thrive by increasing their fitness at the expense of their hosts. Hosts have evolved defense mechanisms, collectively known as immunity, to eradicate parasites or reduce the damage they inflict. The host’s immune system faces a myriad of parasites and is under constant pressure to react quickly and efficiently. Rapid identification and elimination of harmful elements, however, have negative consequences for the host 1,2. Energetic costs associated with establishing an immune system and deploying an immune response have received a lot of attention in evolutionary studies, but only a few have focused on immunopathology and autoimmunity as essential costs of immunity 3–6. When referring to autoimmunity, we mean the damage done by the immune system when it mistakenly identifies self as harmful and then mounts an immune reaction against it. Autoimmune diseases are usually chronic and do not happen necessarily in the presence of parasites. By immunopathology, we refer to the collateral damage inflicted on the host in the course of an immune reaction against a parasite. Immunopathology is, therefore, acute and happens only in the presence of a parasite. For this reason, and following the terminology of classical evolutionary immunology 1,7–9, we classify autoimmunity as constitutive and immunopathology as facultative costs of immunity. When highly sensitive, immune surveillance quickly detects and eliminates parasites, reducing parasite-induced costs. However, increased sensitivity creates two kinds of costs: a lifelong higher probability of mistakenly recognizing self-elements as harmful (autoimmunity) and more collateral damage to host tissues during immune reactions (immunopathology).

There are parallels between the concept of immune sensitivity and that of immune tolerance and resistance 10. Immune resistance refers to host reactions that aim at eradicating the parasite so that the host recovers quickly while immune tolerance refers to reactions aiming at minimizing the costs of infection without eliminating the parasite. Tolerant hosts, therefore, spend more time in the infectious state, which facilitates parasite transmission to other hosts. High immune sensitivity is associated with high immune resistance because it leads to detecting and eliminating parasites rapidly. Low immune sensitivity leads to a higher threshold for deploying immune defenses (see methods, section costs of immune sensitivity) and slower recovery and is thus characteristic of high immune tolerance 11.

Selection on immune sensitivity depends on the balance between the direct damage caused by the parasite, the indirect damage by the immune system during infection (immunopathology), and the lifelong risk of autoimmunity. Consequently, the optimal immune sensitivity depends on the balance between efficient parasite removal and acceptable risks of autoimmunity and immunopathology 5. Metcalf et al. (2017), in one of the first models to study the evolution of immune sensitivity, developed mathematical tools to investigate the evolution of immune sensitivity under a tradeoff between immune sensitivity and specificity. They modeled this tradeoff as two overlapping distributions, one of the host molecular signatures and the other of the parasites. The strength of the tradeoff, consequently, depends on the degree of overlap between the two distributions. They found that the optimal sensitivity increases monotonously with increasing infection hazard (i.e. parasite-induced mortality) and with decreasing mortality due to autoimmunity. These results make intuitive sense: the greater the risk of dying due to infections, the higher is the optimal immune sensitivity. Similarly, the greater the damage due to autoimmunity, the lower should be the immune sensitivity. Albeit an important step, this model lacks two essential elements for studying immunity evolution: an epidemiological framework that captures the ecological feedbacks between immune strategies and the prevalence of infection at the population level and an adaptive dynamics framework to define evolutionary stable strategies.

Previous theoretical studies emphasized the importance of ecological feedbacks to the study of immune strategy evolution and showed how it could change key predictions 4,12–15. Extending these conclusions to the context of immune sensitivity, higher sensitivity leads to shorter infection times, lower infection prevalence, and lower infection rates as a result. This, in turn, decreases the benefits of high sensitivity genes (negative feedback). When genes for high immune specificity spread, hosts remain in the infected state for longer times, the prevalence of the infection increases and selection may favor high specificity (positive feedback) or high sensitivity (negative feedback). Therefore, it is essential to study the evolution of immune sensitivity in an adaptive dynamics framework to find Evolutionary Stable Strategies (ESSs). It is also essential to consider the presence or absence of immune memory for the following reasons. Immunopathology arises only during an infection; therefore, the infection frequency changes the accumulative costs of immunopathology. Autoimmune costs do not depend on the infection frequency and are limited only by the host’s lifespan. In both cases, any given host can get infected multiple times with the same parasite in the absence of immune memory but only once if it possesses lifelong immune memory. Therefore, immune memory may change the balance between the benefits and effective costs of high immune sensitivity differently under acute immunopathological costs versus chronic autoimmune costs. Moreover, when hosts become resistant to infection upon recovery, a higher prevalence of resistant hosts in the host population leads to lower infection prevalence through herd immunity with possible consequences on immune strategies.

Here, we study the evolution of immune sensitivity in the context of an immune discrimination tradeoff under immunopathological or autoimmune costs. We argue that the evolution of sensitivity must be investigated in the context of ecological feedbacks, where infection rates vary according to host strategies and population characteristics. Eco-evolutionary studies usually deal with immune resistance and tolerance as two discrete traits 16,17. We model immune sensitivity as a continuous trait. Hosts can, therefore, either be maximally sensitive, aligning with pure resistance and no tolerance, maximally specific aligning with pure tolerance and no resistance or can fall anywhere in between, displaying different levels of resistance and tolerance. Accounting for the tradeoff between resistance and tolerance, we use adaptive dynamics tools to investigate the following: (i) How does selection act on immune sensitivity under ecological feedbacks between immune strategies and infection prevalence? (ii) How, in that context, do characteristics of the parasite, namely its virulence and degree of host-mimicry, affect immune sensitivity evolution? (iii) How do the lifespan of the host and the physiological nature of immune costs (immunopathological or autoimmune) affect optimal immune sensitivity? For each case, we account for the effect of immune memory on optimal sensitivity and thereby provide predictions on conditions under which selection favors high or low immune sensitivity.

## 2. Results

### 2.1. Optimal sensitivity, parasite virulence and immune tradeoffs

Using classic compartmental models to track the epidemiological dynamics, we applied the nextgeneration matrix approach to find expressions for optimal immune sensitivity under different ecological scenarios 18,19. Considering a resident population at its ecological equilibrium, we derived analytical expressions for the invasion fitness of a rare mutant with slightly different immune sensitivity, assuming that ecological processes occur at a much faster rate than evolutionary change. We found the condition for an Evolutionary Stable Strategy (ESS) of immune sensitivity to be as follows: the ESS host sensitivity maximizes the number of infected hosts at equilibrium. This somewhat counterintuitive finding is analogous to previous results on immune strategy evolution according to which hosts that can sustain a higher parasite population exclude others 1. Therefore, the direction of evolution is determined by the selection gradient 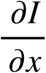 where ***I*** is the number of infected individuals at equilibrium and is the host’s immune sensitivity.

The sign of the selection gradient 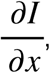, determines the direction of selection on sensitivity. If it is positive, higher sensitivity should be selected. If it is negative, lower sensitivity should evolve. Sensitivity values where 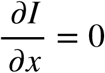 cause the selection gradient to vanish and correspond to possible evolutionarily endpoint.

Considering immunopathological costs, the sign of 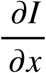 under facultative costs is determined by the signs of the following expressions:

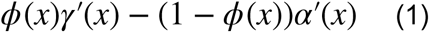

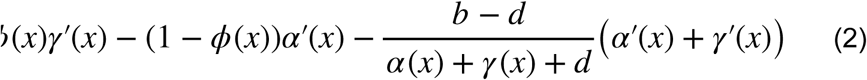

Where (1) and (2) represent situations with or without immune memory respectively. is the death rate due to the infection (the sum of the parasite virulence and extra mortality due to immunopathology as explained in the methods), is the recovery rate, is the *per capita* birth rate of the host, and *d* is the natural *per capita* mortality rate (Table 1). 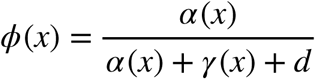 is the probability of death due to infection. The transmission rate *β* does not appear in the derivatives 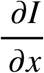, and thus does not affect the ESS immune sensitivity under our assumptions.

**Table 1.** Model parameters and functions

| Parameter and functions | Explanation |
| --- | --- |
| $b$ | *Per capita* birth rate of the host |
| $x$ | Immune sensitivity |
| $q$ | Parameter responds to the similarity between host and parasite molecular signatures. The smaller $q$ is, the higher the overlap between the two. |
| $y(x) = 1 - x^q$ | Immune specificity, function of sensitivity and parasite camouflage. The smaller $q$ is, the stronger the tradeoff between sensitivity and specificity. |
| $d(x) = \mu_b + \mu_i(1 - y(x))$ | Natural death rate of the host (the sum of the host's basal adult mortality rate $\mu_b$ and mortality due to autoimmunity) |
| $\alpha(x) = \alpha_b + \alpha_i(1 - y(x))$ | Death rate due to the infection (the sum of the intrinsic virulence of the parasite $\alpha_b$ and mortality due to immunopathology ) |
| $\gamma(x) = \gamma_b x$ | Recovery rate, function of immune sensitivity and scaling factor $\gamma_b$ |
| $\phi(x) = \frac{\alpha(x)}{\alpha(x) + \gamma(x) + d(x)}$ | The probability of death due to the infection |

Expression (1) predicts that optimal sensitivity depends on the balance between the benefits of increasing sensitivity (i.e. increasing recovery rate *γ*^′^(*x*) = *γ*_b_), weighted by *ϕ*(*x*), and the costs of high sensitivity (i.e. increased mortality during the infection due to higher immunopathology *α*′(*x*) = *qa*_i_*x*^(*q*-1)^ weighted by (1 − *ϕ*(*x*)). Selection favors high immune sensitivity and immune resistance when basal recovery rate γ_b_ and the probability of death due to the infection are high. On the other hand, greater immunopathological costs *α*_i_ expand the range at which lower sensitivity is favored (Supp. Figure 1). Immunopathological costs also depend on *q*, the strength of the tradeoff between sensitivity and specificity. When *q* = 1., the tradeoff between sensitivity and specificity is linear, and selection favors sensitivity values close to zero when the virulence is low or close to one when virulence is high (figure 1 A). As increases, the function defining immune costs becomes accelerating (see methods) and intermediate ESSs exist. The ESS sensitivity increases with higher basal virulence, meaning that infections with more virulent parasites select for higher sensitivity when the costs are immunopathological (Figure 1A).

**Figure 1.**
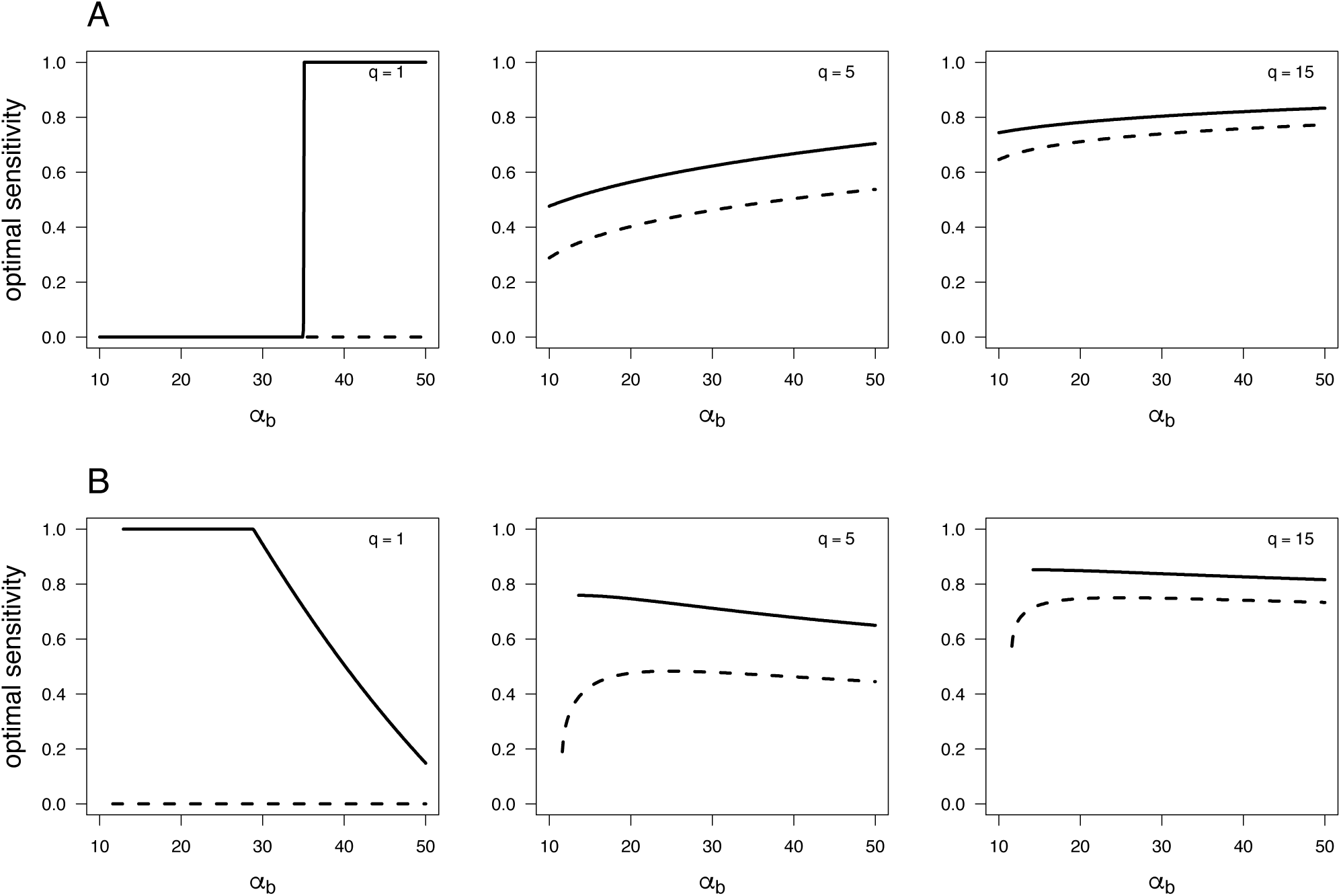
Optimal immune sensitivity (ESS) as parasite virulence (***α*_b_**) is increased, when immune memory is absent (dashed lines) versus when the host possesses immune memory (solid lines). (A) ESS sensitivity when immune costs are acute immunopathology. (B) ESS sensitivity when immune costs are chronic autoimmune. Other parameters values are: in (A) (***μ*_b_ = 3.5, *μ*_i_ = 0, *α*_i_ = 10,*γ* = 1**),and in (B): (***μ*_b_ = 3.5, *μ*_i_ = 1, *α*_i_ = 0,*γ* =** 1).

When the cost of high immune sensitivity is higher autoimmunity (constitutive costs), the natural death rate is a function of immune sensitivity *d*(*x*) (the sum of background mortality and extra mortality due to high sensitivity). The sign of the selection gradient on sensitivity in this case and in the absence of memory is determined by the expression:

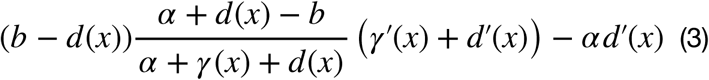

Which contains the benefits and costs of high sensitivity in this scenario. The benefits of high sensitivity (leaving the infected class) are weighted by the *per capita* growth rate (*b* − *d*(*x*))and the probability of death during infection 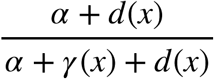 minus the reproductive output during the infection 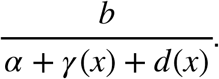. The cost of high sensitivity (death due to autoimmunity) is weighted by the infection-induced mortality, a result that is analogous to Cressler et al. (2015) for selection on recovery rate under constitutive immune costs. When considering autoimmune costs in the presence of immune memory, the expression for 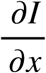 becomes analytically intractable (supplementary table 1) but EES immune sensitivity can still be numerically found.

Under autoimmune costs, intrinsic parasite virulence *α*_b_ has an opposite effect on ESS sensitivity with maximal optimal sensitivity at low rather than high parasite virulence (Figure 1). To understand this result, it is first important to remember that immunopathological costs are suffered by infected hosts only, whereas autoimmune costs are paid by hosts in all infection states. Even when the probability of getting infected is low, it always pays to be highly sensitive under immunopathological costs because the host suffers these costs only when infected. Under autoimmune costs, hosts with high immune sensitivity may have an advantage when infected with a virulent parasite but suffer more lifelong autoimmunity. This means that, under autoimmune costs, the benefits of high sensitivity are limited by the probability and length of infection while the costs are limited only by the lifespan. Therefore, high virulence may then select for lower sensitivity (Figure 1 B) because of the epidemiological effect of high virulence on infection rate: when parasites are very virulent, infected hosts die at higher rates and susceptible hosts have a *de facto* lower probability of contracting the disease. This effect of increasing virulence is also mathematically demonstrated in expression (3) as follows. The first term in (3) is weighted by 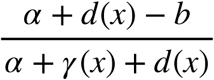 which is a saturating increasing function of *α*, so higher virulence may select for higher sensitivity. However, this effect is counterbalanced by the second term *αd*′(*x*). Since the weight of the first term is a saturating function of *α*(*x*), increasing *α*(*x*) beyond a certain point will increase *α*(*x*)*d*′(*x*) more than 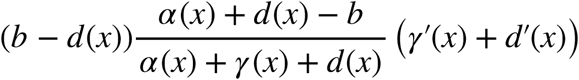 and weaken selection for high sensitivity.

### 2.2. The effect of host characteristics on optimal immune sensitivity

Equation (1) shows that increased natural mortality selects for lower optimal immune sensitivity under immunopathological costs and immune memory since *ϕ*(*x*) is a decreasing function of natural mortality rate *d*. Equation (2) predicts optimal sensitivity in hosts without immune memory and differs from equation (1) only by its last term representing the sum of both derivatives (*α*′(*x*)+*γ*′(*x*)) weighted by the *per capita* growth rate *b − d* and the duration of the infection 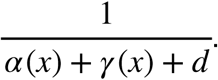. Since we have assumed that the natural birth rate is greater than the natural mortality rate (*b − d* > 0, which is required for the stability of the epidemiological equilibrium as explained in the methods), equation (2) is negative for a larger range of parameter values than equation (1). This means that under immunopathological costs, and for any set of parameter values, ESS sensitivity is always lower when hosts lack immune memory compared to when they become resistant to infections after recovery. The difference in optimal immune sensitivity between hosts with or without immune memory is proportional to the host *per capita* growth rate *b − d*. This prediction is biologically intuitive if the host suffers immune costs during the infection only. A host that possesses immune memory will suffer these costs only once in her lifetime because she remains immune to the pathogen if she survives the infection. In contrast, without immune memory, hosts can be infected multiple times, which means greater lifetime cumulative immunopathological costs. When *d* is high, the expressions for optimal sensitivity with or without immune memory tend to the same value for the following reason. When the death rate due to extrinsic factors that are independent of infection is high, recovered immune hosts rapidly disappear from the population, eliminating the epidemiological effect of immune memory. Therefore, optimal immune sensitivity tends towards the same value for hosts with or without immune memory when extrinsic mortality is high (Figure 2, A). In hosts lacking immune memory and under immunopathological costs, optimal immune sensitivity, therefore, increases with increasing ***μ*_b_** (Figure 2, A) and this is consistent with previous model predictions made without accounting for epidemiological feedbacks 6. When hosts possess immune memory, however, the contrary is true and hosts with longer lifespans evolve higher immune sensitivity.

**Figure 2.**
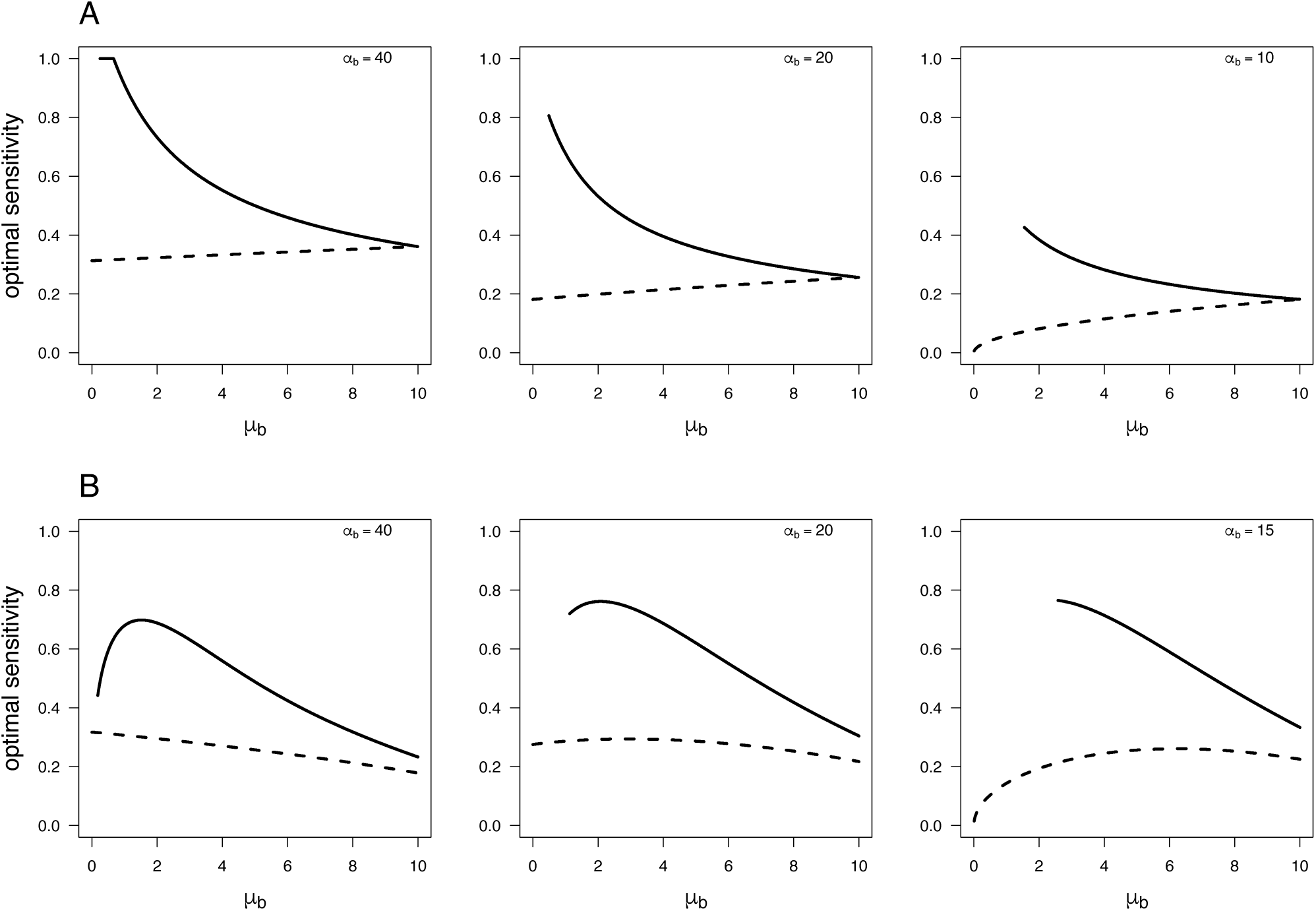
Optimal immune sensitivity as host basal mortality rate (***μ*_b_**) is increased, when immune memory is absent (dashed lines) or when the host possesses immune memory (solid lines). (A) ESS sensitivity when immune costs are acute immunopathology (facultative). (B) ESS sensitivity when immune costs are chronic autoimmune (constitutive). Other parameters values are: in (A) (***α*_b_ = 20, *μ*_i_ = 0, *α*_i_ = 10, *γ*_b_ = 1**),and in (B): (***α*_b_ = 20, *μ*_i_ =, *α*_i_ = 0, *γ*_b_ = 1**).

Selection on optimal immune sensitivity follows a different pattern under autoimmune costs. Autoimmunity is a type of constitutive cost and is suffered throughout life, such that hosts with longer basal lifespan (i.e., lower ***μ*_b_**) have a greater chance of dying from autoimmunity. This also means that hosts with potentially longer lifespans must balance the risk of dying from an infection with the risk of autoimmunity. Looking at hosts lacking immune memory (dashed curves in Figure 2B), it seems that the balance tips over in favor of greater sensitivity for long-lived hosts. Only when parasite virulence is extremely low is it advantageous to have low sensitivity when lifespan is long (rightmost graph in Figure 2B). When hosts do possess immune memory, recovered individuals become abundant under longer lifespans. This decreases infection rates through a process similar to herd immunity. It then pays to be less immune sensitive to avoid autoimmune costs (leftmost graph in Figure 2B). When basal mortality increases (natural lifespan decreases), resistant individuals die rapidly, resulting in greater infection rates and selecting for higher immune sensitivity. When basal mortality increases further, epidemiological feedbacks lead to a relative decrease in infection rate when the population turnover is high enough (similar to the case of facultative costs above) and it is then advantageous to have lower immune sensitivity again (Figure 2B). Therefore, host species with intermediate lifespan are expected to have the highest immune sensitivity in this case scenario (autoimmune costs in hosts that possess long-term acquired immune memory).

## 3. Discussion

Selection acts on immune sensitivity to balance the risk of damaging self with sufficient protection against parasites. While previous models have found that the virulence of the parasite and life-history traits of the host can monotonously affect immune sensitivity evolution, this is the first study, that combines immune discrimination tradeoffs with ecological feedbacks. We modeled optimal immune sensitivity under a biologically meaningful immune discrimination tradeoff and showed how the physiological cost structure and the presence of immune memory affect immune sensitivity evolution in response to parasite virulence and host lifespan. We compare our results to models on immune sensitivity evolution before discussing them in a more general context of host immunity evolution.

Our first result is that increasing parasite virulence selects immune responses to be more intense but does not increase risk of chronic autoimmunity. One previous model had examined the effect of parasite virulence on host sensitivity but did not account for the effect of host strategy on infection prevalence, and hence the ecological feedback between the two 6. Hardly surprising, they have found a monotonous effect of virulence, where higher parasite-induced mortality selects for higher optimal sensitivity. Even though their result appears intuitive, the effect of parasite virulence on immune sensitivity evolution can be more nuanced. We show here that the effect of parasite virulence qualitatively differs when the costs are facultative versus when they are constitutive. Immune memory does not qualitatively change the effect of parasite virulence on optimal sensitivity even though sensitivity is always higher when the host possesses immune memory. The pattern found by Metcalf et al. (2017), therefore, cannot be generalized to the autoimmune cost scenario. The evolutionarily optimal sensitivity depends on the balance between the benefits and costs of increasing sensitivity. When the costs are immunopathological, they incur during the infection only and thus depend on the length and frequency of infections. When the cost of high sensitivity is an increased risk of autoimmunity, the costs are constant life long. A highly virulent parasite might be deadly for those who get it, but the probability of encountering it is lower due to the epidemiological effect of virulence. Therefore, when costs are autoimmune, high virulence reduces the effective benefits of high sensitivity but not the costs and may select for lower immune sensitivity. More general models on the role of parasite virulence in the evolution of immune strategies have found contradictory results 1,2,7,17,20,21. We show here that the response of the host optimal immunity to changes in parasite virulence depends on three factors: the physiological cost nature (facultative or constitutive), the presence of immune memory, and the shape of the tradeoff that depends, in our model, on the overlap between host and parasite molecular signatures. Both Day et al. (2003) and Boots et al. (2018) 2,7, for example, found that high parasite virulence always selects for higher induced defenses without accounting for immune memory. We confirm this result here and show that this effect is even stronger in the presence of immune memory. Van Baalen (1998), in a co-evolutionary model, examined the effect of virulence on immunity evolution and found that it pays to invest in an immune system only at intermediate values of parasite virulence22. This result is echoed in our findings where the costs are constitutive and the host does not possess acquired immune memory, which are also the assumptions of this model. Restif and Koella (2004) studied the evolution of resistance and tolerance as two independent traits and examined different possible functions for the costs of immune investment (linear or quadratic, additive or multiplicative). They found that the effect of virulence can be monotonic or u-shaped depending on the shape of the cost function. We show the same effect by combining the effect of *q* (tradeoff strength) and virulence, where *q* changes the range where virulence is associated with increased or decreased sensitivity.

The effect of host lifespan depends on the nature of costs and the presence of immune memory. This differs from previous findings on immune sensitivity evolution and results from our inclusion of ecological feedbacks and immune memory. When we consider selection due to immunopathological costs (inflammation, fever, etc.), we expect short-lived animals to have higher immune sensitivity (the dashed lines on figure 2A). Short-lived species apparently care less about the accumulation of collateral damage due to their immune responses. This pattern is, however, reversed if recovery leads to lifelong acquired memory (solid line on the figure). In this case, the host can get infected only once in a lifetime. This means no accumulative damage to be afraid of and the longer the host lives, the more it will benefit from immune protection. This pattern is also different when we consider the lifelong higher risk of developing autoimmune diseases due to a higher chance of false positives associated with high sensitivity. In this case, and in the absence of immune memory, the shorter a host lives, the lower should be its immune sensitivity. The reason behind this is the way we have incorporated autoimmunity into our model. We have added an extra mortality term that resembles autoimmunity, and the probability of suffering this extra term is directly related to immune sensitivity. Species whose lifespans are already short cannot afford to shorten it further for extra immune protection. Nonetheless, the slope is not very steep because of the complicated effect of mortality on infection risk, population turnover, etc. When the host possesses immune memory, there is an extra phenomenon of hosts with lifelong spans evolving low immune sensitivity (hump-shaped solid line in 2B). That is because of the effect of herd immunity and resistant hosts remaining for a longer time in the population, reducing the risk of contracting the infection.

Our results may provide evolutionary explanations for some observations related to immune sensitivity. For example, we have shown that higher parasite virulence selects for higher immune sensitivity in induced immune responses (characterized by a high degree of immunopathology). This makes highly virulent parasites double problematic since they combine high intrinsic virulence with selection on the host for high immunopathology, leading to a high Case Fatality Ratio (CFR). This may partially explain phenomena like the cytokine storm, an uncontrolled over-production of soluble markers of inflammation producing an uncontrolled and generalized inflammatory response, particularly by the innate immune system 23. The cytokine storm has been associated with viruses known to be highly virulent like H5N1 influenza 24, dengue hemorrhagic-fever 23, acute respiratory distress syndrome (ARDS) caused by SARS coronavirus (SARS-CoV) MERS coronavirus (MERS-CoV) 25, and even the H1N1 Spanish flu 26. Moreover, most of these viruses are newly emergent without enough time to specialize on human hosts to avoid immune detection through host mimicry. This corresponds to an increasing *q* in our model, which extends the range of virulence values associated with high immune sensitivity (Supp. Figure S1).

Low immune sensitivity is always favored when the intrinsic virulence is low (low immune sensitivity when the costs are immunopathological). When the intrinsic virulence is high, low immune sensitivity can still evolve if the costs are autoimmune possibly resulting in chronic infections. Moreover, the greater the overlap between self-antigens and those of the microorganisms, the greater the self-damage in the course of an immune reaction, and the stronger is the selection for immune tolerance. Therefore, the presence of microorganisms with high degrees of mimicry could favor the evolution of immune tolerance (low immune sensitivity)27. This could be the case of parasites that highly specialize in immune evasion of their host. One possible example is *Mycobacterium tuberculosis*. *Mycobacterium tuberculosis* (Mtb) has coevolved with humans for 70,000 years 28. Unlike the opportunistically parasitic and the non-pathogenic soil dwellers species of mycobacteria, Mtb has no known environmental reservoirs and does not survive outside human hosts pointing to its highly specialized nature 29. Interestingly, 90–95% of infected individuals (9 out of 10 people) are resilient or tolerant to the presence of Mtb without any disease symptoms 30. Our results indicate that the highly specialized nature of Mtb and the potentially high immunopathological costs associated with eradicating it could have selected immune responses to be tolerant to it.

Future studies should extend our framework to include other possible mechanisms of tolerance that might be independent of or even positively correlated with resistance. We have focused on low immune sensitivity as one mechanism of immune tolerance. When hosts evolve low immune sensitivity, their immune system is more likely to „ignore“ the parasite, sparing the host immunopathological costs of resistance without decreasing the fitness of the parasite. This falls under the definition of immune tolerance 10 but does not include all its possible mechanisms. Increasing tissue repair, for example, is another way to tolerate parasites and does not involve immune sensitivity (e.g., increasing red cell production in the course of malaria infection). Moreover, we have taken a host-centric viewpoint. In natural systems, hosts and parasites co-evolve. Tolerance could select for more virulent strains when virulence is correlated with a higher transmission rate 14 and the evolved high virulence may select for less tolerance (immunopathological costs) or more tolerance (due to autoimmune costs). This might affect the evolutionary trajectory of virulence when it encompasses parasite-induced mortality and selection on the host immune sensitivity and requires co-evolutionary models. Co-evolutionary models should also consider the concurrent evolution of the parasite mimicry and the host discrimination power to counter mimicry. Moreover, it is possible to test our predictions experimentally. For example, one could investigate selection on genes that cause immune reactions to be more intense (immunopathology high sensitivity) versus genes that increase the risk of developing autoimmune diseases (autoimmune high sensitivity) and how they respond to changes in parasite virulence or the lifespan of the host, in model oragnisms possessing or lacking immune memory. We have not accounted for a resource-reproduction tradeoff in our model. The reason is that many models examined the economics of immunity and particularly dividing resources between immunity and reproduction. Ours is the first to examine the evolution of immune sensitivity at the population level and distinguish between immunopathological and autoimmune costs. We wanted to keep it simple at this stage and focus on how these two types of cost can drive the evolution of immunity in different directions.

This is the first model to our knowledge to combine demographically framed tradeoffs between immune sensitivity and specificity with epidemiological feedbacks, immune physiological cost structure (immunopathological vs autoimmune), and acquired immune memory. We have shown that the effect of parasite virulence depends on the physiological nature of costs while the effect of host lifespan depends on the presence or absence of immune memory. Our approach is general and can be broadly applied to understanding the evolution of immune responses across a broad range of the tree of life.

## 4. Methods

### 4.1. Epidemiological models

We based our study on compartmental models of parasite transmission. We used an SIS model (Susceptible-Infected-Susceptible) for the case of hosts with no immune memory (figure 3) and an SIR model (Susceptible-Infected-Recovered) for the case of long-term immune memory (figure 4). The two models were built using the set of ordinary differential equations presented in Table 2.

**Table 2.** ODEs defining the SIS and SIR models

| No immune memory | Immune memory |
| --- | --- |
| $\frac{dS}{dt} = b(S + I) - \beta SI - dS + \gamma I$ (4) | $\frac{dS}{dt} = b(S + I) - \beta SI - dS$ (6) |
| $\frac{dI}{dt} = \beta SI - (\alpha + \gamma + d)I$ (5) | $\frac{dI}{dt} = \beta SI - (\alpha + \gamma + d)I$ (7) |
| | $\frac{dR}{dt} = \gamma I - dR$ (8) |

**Figure 3.**
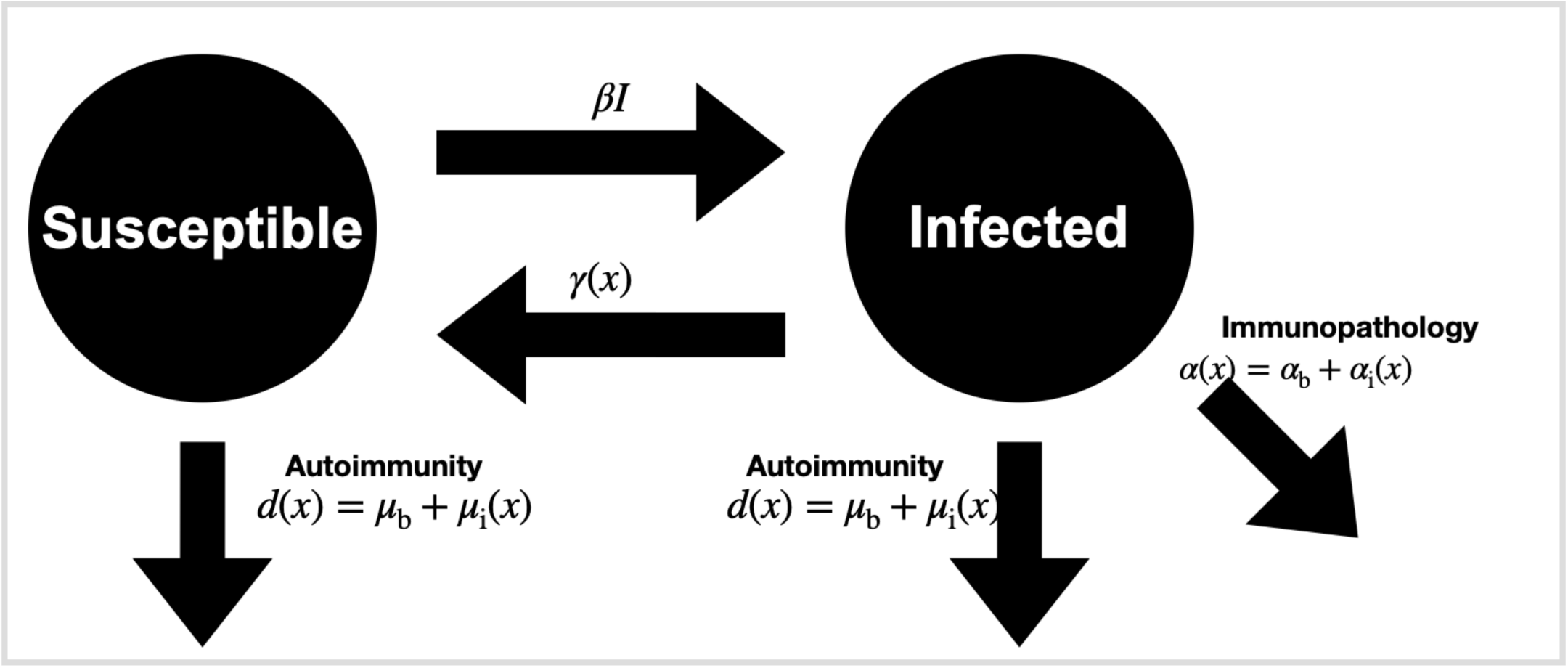
Schematic representation of the ecological feedbacks in the SIS epidemiological model when the host lacks acquired immune memory. Hosts can get infected at a rate *βI*, proportional to the frequency of infected individuals. Infected hosts can recover at rate ***γ* (*x*)**, a function of their immune sensitivity. Once infected, hosts die at mortality rate ***α*** that is the sum of parasite virulence and immunopathology which is also a function of sensitivity ***x***. Hosts in both classes suffer natural mortality ***d***, the sum of background mortality and autoimmune mortality, also a function of immune sensitivity.

**Figure 4.**
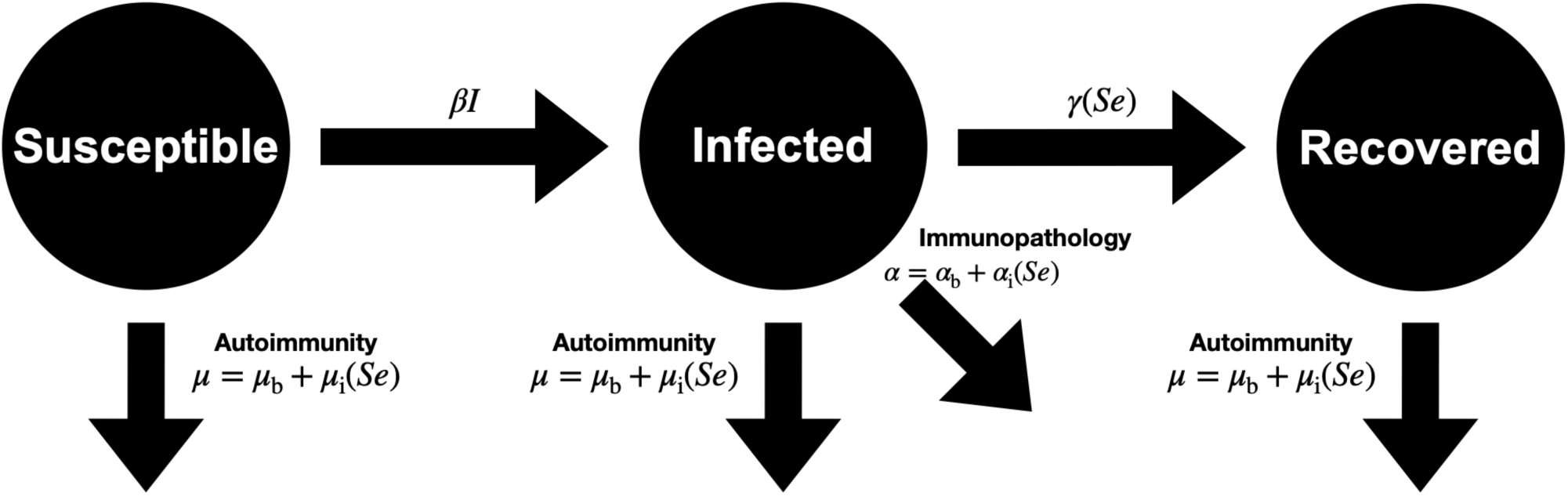
Schematic representation of the ecological feedbacks in the SIR epidemiological model when the host possesses acquired immune memory. Once recovered, the hosts becomes resistant to re-infection. Other assumptions remain the same as in the SIS model

**Figure 5.**
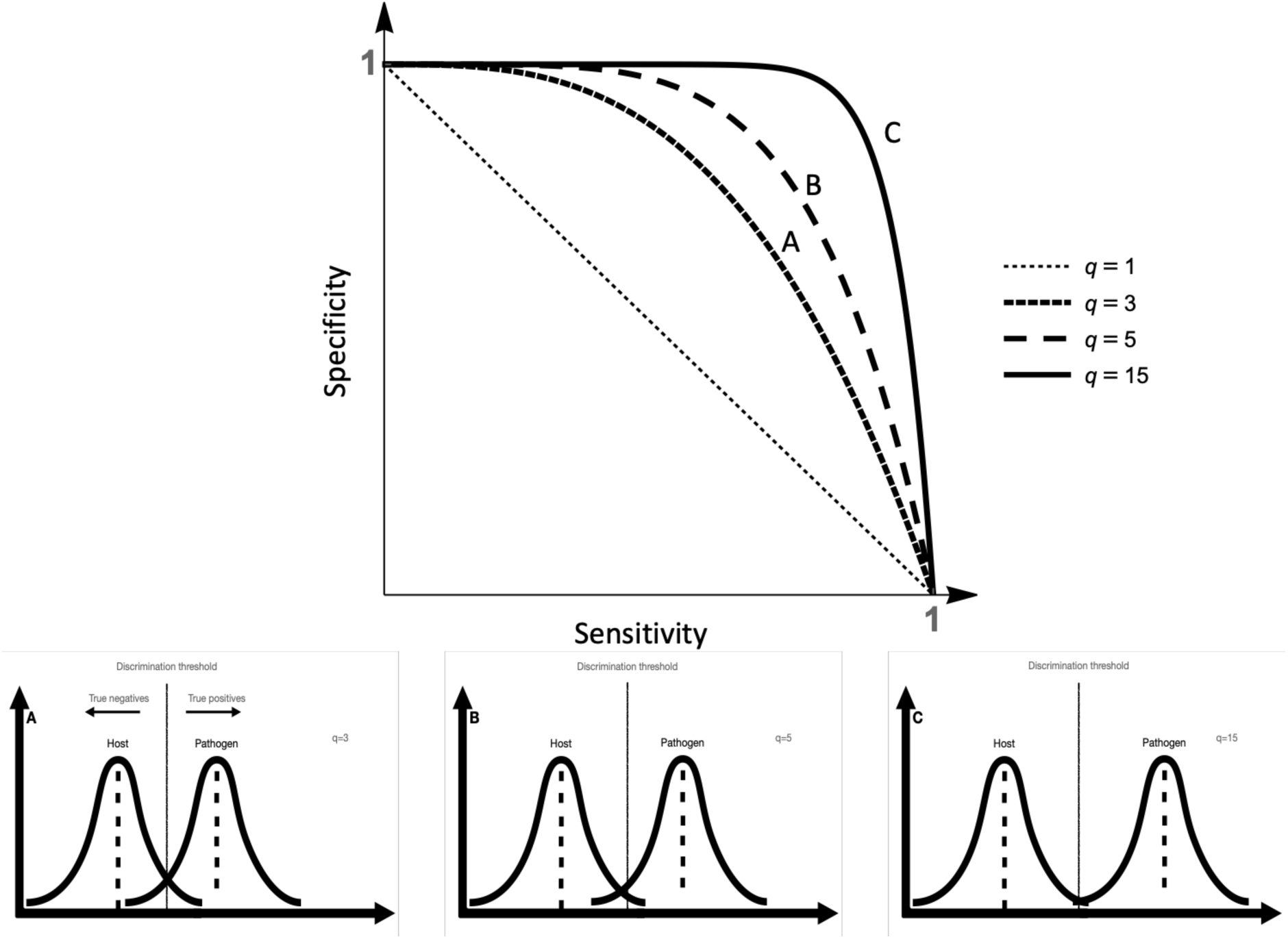
Schematic depiction of the immune discrimination tradeoff and the relationship between immune sensitivity and specificity. The figures at the bottom represent the distributions of the host and pathogen molecular signatures at three degrees of overlap. The figure on the top represents the corresponding sensitivity-specificity tradeoff line. The further apart the two distributions are, the higher is the discrimination resolution and the weaker is the immune tradeoff

In these equations, S, I, and R are the numbers of susceptible, infected, and Recovered hosts respectively. The remaining parameters are explained in Table 1 in the result section. Throughout, we assume that *α + d > b > d*. This means that the host has a negative *per capita* growth rate when infected and a positive one when uninfected. This is required for the stability of the epidemiological equilibrium; the population would grow infinitely in the absence of the parasite but parasite related mortality keeps it in check.

To calculate optimal immune sensitivity, we assumed a homogenous resident host population at demographic equilibrium. To find the expressions for the number of hosts in each class at epidemiological equilibrium, we solve equations (4) to (8) (table 2) equal to zero for S, I, and R to arrive at the following expressions for the number of individuals in each class at equilibrium (Table 3).

**Table 3.** Numbers of Susceptible, Infected, and Recovered hosts at equilibrium

| No immune memory | Immune memory |
| --- | --- |
| $S = \frac{\alpha + \gamma + d}{\beta} \quad (9)$ | $S = \frac{\alpha + \gamma + d}{\beta} \quad (11)$ |
| $I = \frac{(b - d)(\alpha + \gamma + d)}{\beta(\alpha - (b - d))} \quad (10)$ | $I = \frac{(b - d)d(\alpha + \gamma + d)}{\beta(d(\alpha + \gamma + d) - b(\gamma + d))} \quad (12)$ |
| | $R = \frac{(b - d)\gamma(\alpha + \gamma + d)}{\beta(d(\alpha + \gamma + d) - b(\gamma + d))} \quad (13)$ |

### 4.2. Costs and benefits of immune sensitivity

We assume that is the host’s immune sensitivity, where 1 > *x* > 0. By increasing immune sensitivity, the host’s recovery rate increases such that:

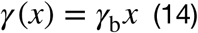

Where *γ*_*b*_, the basal recovery rate, is a scaling factor. Therefore, hosts have the maximal possible recovery rate when sensitivity equals one, and do not recover at all if sensitivity is equal to zero. Immune surveillance faces two overlapping molecular distributions: one belonging to the parasite and another of the host elements. The immune system has to discriminate between these two groups efficiently; however, it is still prone to error. This problem is exacerbated by molecular mimicry, the selection on pathogens to resemble harmless elements to evade detection 31. In that context, sensitivity and specificity are statistical measures of discrimination accuracy. Sensitivity measures the probability of classifying a true positive as such (here, the proportion of parasitic elements that are correctly detected by the immune system) and is closely related to the concept of type II error in statistical testing (i.e. high sensitivity reduces the probability of accepting the null hypothesis when the alternative hypothesis is true). Specificity measures the probability of classifying true negatives as such (here, the proportion of harmless elements that are correctly classified as such) and is related to the concept of type I error in hypothesis testing (e.g. high specificity reduces the probability of rejecting the null hypothesis when it is true) 32. The relationship between sensitivity and specificity accordingly depends on the degree of overlap between the parasite and host antigen distributions 33. If we assume that *q* corresponds to the distance between the distribution means of the host’s and parasite’s molecular signatures, the relationship between sensitivity *y* and specificity *x* can be captured by:

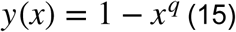

When sensitivity is maximal (*x* = 1) specificity equals zero and when sensitivity is zero, specificity equals one. At intermediate values of sensitivity, specificity depends on *q* (Figure 1). When *q* = 1, the relationship between sensitivity and specificity is linear corresponding to completely overlapping distributions of the host and parasite antigens, leading to equal chances of true negatives and positives. When *q* increases, the resolution of immune discrimination increases, decreasing the strength of the tradeoff between sensitivity and specificity and allowing greater increases in sensitivity without big decreases of specificity.

We considered two types of costs related to decreasing immune specificity: facultative costs (immunopathology) suffered by infected individuals only (due to inflammation, fever, etc..) whereas constitutive costs (autoimmunity) can be suffered by hosts in all classes (due to mistakenly recognising self as parasitic). To model facultative costs we assume that the mortality rate due the infection is the sum of *α*_b_, the intrinsic virulence of the parasite (direct damage to the host) and *α*_i_, the indirect collateral damage done to the host tissues due to its own immune response (i.e. attacking oneself through false positive parasite identification). The mortality rate due to infection is given by the following equation:

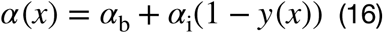

Where *y* resembles the immune specificity. The relationship between immune sensitivity and specificity is determined by equation (15). When *y* equals one, immune responses are completely specific, and hosts suffer only direct parasite-induced mortality without any collateral damage resulting from false positives. When specificity equals zero, facultative costs are maximal and infected hosts suffer maximal possible infection mortality rate (*α*(*y*) = *α*_b_ + *α*_i_).

To model constitutive immune costs, we assumed that immune sensitivity affects the host’s natural mortality rate *d*, affecting all host classes. Similar to the facultative costs case,*d* is the sum of the background mortality *μ*_b_, and the autoimmune-mortality due to false positives and immune reaction against self:

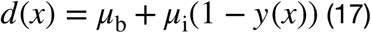

When immune surveillance is perfect, equals one and the average lifespan of an uninfected host equals 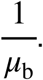. When specificity equals zero, the host suffers maximal natural mortality and the average lifespan of uninfected hosts equals 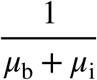.

Our definitions of costs and benefits of increasing immune sensitivity or specificity align with the concepts of immune resistance and tolerance 10. By increasing immune sensitivity, the host increases its recovery rate *γ* but also increases the infection-related mortality rate *α* (facultative cost) or the natural mortality rate *d* (constitutive cost) as a result of decreased immune specificity. Increasing immune sensitivity decreases the time the host spends in the infected class and decreases the parasite fitness which constitutes parasite resistance. Increasing specificity decreases the fitness loss due to the infection (by lowering facultative and constitutive costs) but increases the parasite fitness (by decreasing host recovery rate), which is a type of parasite tolerance. The shape of the relationship between immune sensitivity and specificity is presented in Box 1. Briefly, parameter *q* (corresponding to the resolution of immune discrimination) determines the shape of the tradeoff between host resistance and tolerance, such that lower *q* leads to a stronger tradeoff (Box 1).

### 4.3. Invasion analysis

To calculate evolutionary stable strategies (ESS) for immune sensitivity, we study the conditions for the invasion of a rare mutant in a resident population at its endemic equilibrium. We assume that the prevalence of the mutant is so low that its epidemiological effect can be ignored (e.g. it does not affect the infection probability of individuals in the resident population). To arrive at the expressions for invasion fitness, we use the Next Generation Matrices Tools 18,19. The following local stability matrices (i.e. Jacobean matrices) are used to determine whether or not a rare mutant invades a population of resident in situations where hosts lack or possess immune memory respectively:

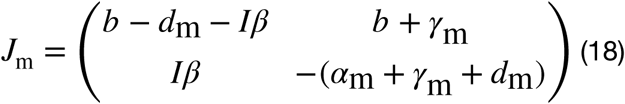

For the SIS model, and:

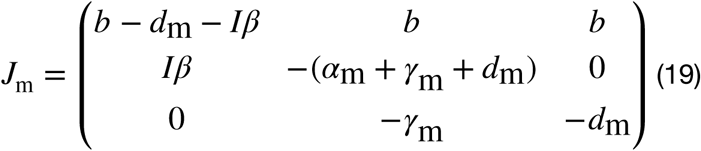

For the SIR model.

We then decompose *J*_m_ into an inflow matrix *F* and an outflow matrix *V* for (18) and (19) separately, such that *V* is nonsingular, *F* and *V*^−1^ are non-negative, and the real parts of all eigenvalues of −*V* are negative. If we define as the spectral radius of a matrix (the maximum absolute value of all the eigenvalues), it can be shown that a rare mutant will invade if *ρ*(*FV*^−1^) > 1 18, meaning that:

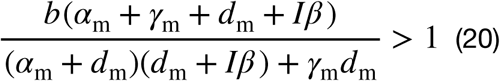

for the SIS model and:

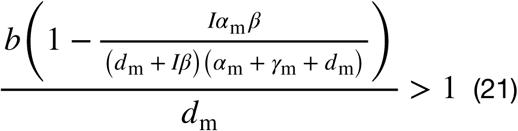

for the SIR model.

Rewriting (19) and (20) by separating *I* (the resident infectious class equilibrium) from the mutant terms, we get:

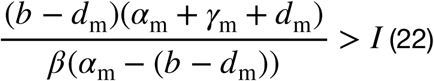

For the SIS model. The expression on the left of (22) is the size of the infected class at the equilibrium set by the mutant. Similarly, in the SIR model, invasion requires that:

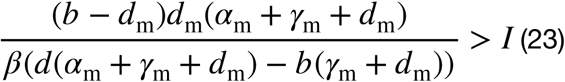

The expression on the left is analogously the size of the infected class set by the mutant (compare (23) and (12)). Analogous to results in community ecology, evolution therefore maximizes *I*_m_; it favors the host that can sustain the highest parasite population.

## Author contributions

L.A., M.G., N.B. and F.R. conceived the project. L.A. designed and performed the mathematical modeling and wrote the first draft with the help and support from M.G.; all authors revised and edited the manuscript.

## Acknowledgments

We thank Hanna Kokko for helpful comments and Abagail Breidenstein for proofreading. The Mäxi Foundation, Zurich, funded the project through a grant to F.R.

## Literature

1 Cressler, C. E., Graham, A. L. & Day, T. Evolution of hosts paying manifold costs of defence. Proceedings of the Royal Society B-Biological Sciences 282, doi:ARTN 20150065 10.1098/rspb.2015.0065 (2015).

2 Day, T. & Burns, J. G. A consideration of patterns of virulence arising from host-parasite coevolution. Evolution 57, 671–676 (2003).

3 Boots, M. & Bowers, R. G. The evolution of resistance through costly acquired immunity. Proceedings of the Royal Society B-Biological Sciences 271, 715–723, doi:10.1098/rspb.2003.2655 (2004).

4 van Boven, M. & Weissing, F. J. The evolutionary economics of immunity. American Naturalist 163, 277–294, doi:10.1086/381407 (2004).

5 Graham, A. L., Allen, J. E. & Read, A. F. Evolutionary causes and consequences of immunopathology. Annual Review of Ecology Evolution and Systematics 36, 373–397, doi:10.1146/annurev.ecolsys.36.102003.152622 (2005).

6 Metcalf, C., Tate, A. & Graham, A. Demographically framing trade-offs between sensitivity and specificity illuminates selection on immunity. Nature Ecology & Evolution 1, 1766–1772, doi:10.1038/s41559-017-0315-3 (2017).

7 Boots, M. & Best, A. The evolution of constitutive and induced defences to infectious disease. Proc Biol Sci 285, doi:10.1098/rspb.2018.0658 (2018).

8 Hamilton, R., Siva-Jothy, M. & Boots, M. Two arms are better than one: parasite variation leads to combined inducible and constitutive innate immune responses. Proceedings of the Royal Society B-Biological Sciences 275, 937–945, doi:10.1098/rspb.2007.1574 (2008).

9 Shudo, E. & Iwasa, Y. Inducible defense against pathogens and parasites: Optimal choice among multiple options. Journal of Theoretical Biology 209, 233–247, doi:10.1006/jtbi.2000.2259 (2001).

10 Raberg, L., Graham, A. L. & Read, A. F. Decomposing health: tolerance and resistance to parasites in animals. Philosophical Transactions of the Royal Society B-Biological Sciences 364, 37–49, doi:10.1098/rstb.2008.0184 (2009).

11 Medzhitov, R., Schneider, D. S. & Soares, M. P. Disease Tolerance as a Defense Strategy. Science 335, 936–941, doi:10.1126/science.1214935 (2012).

12 Boots, M., Best, A., Miller, M. R. & White, A. The role of ecological feedbacks in the evolution of host defence: what does theory tell us? Philosophical Transactions of the Royal Society B-Biological Sciences 364, 27–36, doi:10.1098/rstb.2008.0160 (2009).

13 Donnelly, R., White, A. & Boots, M. The epidemiological feedbacks critical to the evolution of host immunity. J Evol Biol 28, 2042–2053, doi:10.1111/jeb.12719 (2015).

14 Restif, O. & Koella, J. C. Shared control of epidemiological traits in a coevolutionary model of host-parasite interactions. American Naturalist 161, 827–836, doi:Doi 10.1086/375171 (2003).

15 Roy, B. A. & Kirchner, J. W. Evolutionary dynamics of pathogen resistance and tolerance. Evolution 54, 51–63, doi:10.1111/j.0014-3820.2000.tb00007.x (2000).

16 Boots, M. & Bowers, R. G. Three mechanisms of host resistance to microparasites - Avoidance, recovery and tolerance - Show different evolutionary dynamics. Journal of Theoretical Biology 201, 13–23, doi:Doi 10.1006/jtbi.1999.1009 (1999).

17 Restif, O. & Koella, J. C. Concurrent evolution of resistance and tolerance to pathogens. American Naturalist 164, E90–E102, doi:Doi 10.1086/423713 (2004).

18 Diekmann, O., Heesterbeek, J. A. P. & Roberts, M. G. The construction of next-generation matrices for compartmental epidemic models. Journal of the Royal Society Interface 7, 873–885, doi:10.1098/rsif.2009.0386 (2010).

19 Hurford, A., Cownden, D. & Day, T. Next-generation tools for evolutionary invasion analyses. Journal of the Royal Society Interface 7, 561–571, doi:10.1098/rsif.2009.0448 (2010).

20 Boots, M. & Haraguchi, Y. The evolution of costly resistance in host-parasite systems. American Naturalist 153, 359–370, doi:Doi 10.1086/303181 (1999).

21 Carval, D. & Ferriere, R. A Unified Model for the Coevolution of Resistance, Tolerance, and Virulence. Evolution 64, 2988–3009, doi:10.1111/j.1558-5646.2010.01035.x (2010).

22 van Baalen, M. Coevolution of recovery ability and virulence. Proc Biol Sci 265, 317–325, doi:10.1098/rspb.1998.0298 (1998).

23 Tisoncik, J. R. et al. Into the Eye of the Cytokine Storm. Microbiology and Molecular Biology Reviews 76, 16–32, doi:10.1128/Mmbr.05015-11 (2012).

24 Yuen, K. Y. & Wong, S. S. Human infection by avian influenza A H5N1. Hong Kong Med J 11, 189–199 (2005).

25 Channappanavar, R. & Perlman, S. Pathogenic human coronavirus infections: causes and consequences of cytokine storm and immunopathology. Semin Immunopathol 39, 529–539, doi:10.1007/s00281-017-0629-x (2017).

26 Liu, Q., Zhou, Y. H. & Yang, Z. Q. The cytokine storm of severe influenza and development of immunomodulatory therapy. Cell Mol Immunol 13, 3–10, doi:10.1038/cmi.2015.74 (2016).

27 Rook, G. A. W. et al. Mycobacteria and other environmental organisms as immunomodulators for immunoregulatory disorders. Springer Semin Immun 25, 237–255, doi:10.1007/s00281-003-0148-9 (2004).

28 Gutierrez, M. C. et al. Ancient origin and gene mosaicism of the progenitor of Mycobacterium tuberculosis. Plos Pathogens 1, 55–61, doi:ARTN e5 10.1371/journal.ppat.0010005 (2005).

29 Saelens, J. W., Viswanathan, G. & Tobin, D. M. Mycobacterial Evolution Intersects With Host Tolerance. Front Immunol 10, doi:ARTN 528 10.3389/fimmu.2019.00528 (2019).

30 Divangahi, M., Khan, N. & Kaufmann, E. Beyond Killing Mycobacterium tuberculosis: Disease Tolerance. Front Immunol 9, doi:ARTN 2976 10.3389/fimmu.2018.02976 (2018).

31 Hurford, A. & Day, T. Immune Evasion and the Evolution of Molecular Mimicry in Parasites. Evolution 67, 2889–2904, doi:10.1111/evo.12171 (2013).

32 Loong, T. W. Understanding sensitivity and specificity with the right side of the brain. Bmj-Brit Med J 327, 716–719, doi:Doi 10.1136/bmj.327.7417.716 (2003).

33 Gaddis, G. M. & Gaddis, M. L. Introduction to biostatistics: Part 3, Sensitivity, specificity, predictive value, and hypothesis testing. Ann Emerg Med 19, 591–597, doi:10.1016/s0196-0644(05)82198-5 (1990).

