## Supplemental table 1 for "The Evolution of Immune Sensitivity under Immunopathological and Autoimmune Costs"

Supplementary materials

### 1. Analytical results tables:

Supplementary Table 1. Expressions for the derivative that determines the selection gradient on immune sensitivity when the costs are immunopathological or autoimmune and when the host possesses or lacks acquired immune memory

| Cost type | No Memory | Memory |
|---|---|---|
| **Immunopathology** | $\phi(x)\gamma'(x) - (1-\phi(x))\alpha'(x) - \frac{b-d}{\alpha(x)+\gamma(x)+d}(\alpha'(x)+\gamma'(x))$ | $\phi(x)\gamma'(x) - (1-\phi(x))\alpha'(x)$ |
| **Autoimmunity** | $(b-d(x))\frac{\alpha(x)+d(x)-b}{\alpha(x)+\gamma(x)+d(x)}\left(\gamma'(x)+d'(x)\right) - \alpha(x)d'(x)$ | $\alpha b\left((b-d)d\left(\gamma'+d'\right)\right) - \left(((\alpha+\gamma+d)(b-d)-\alpha b)^2 - (\alpha b)^2 + \alpha b^2(\alpha+\gamma+d)\right)d'$ |

Supplementary Table 2. Model parameters and functions

| Parameter and functions | Explanation |
|---|---|
| $b$ | *Per capita* birth rate of the host |
| $x$ | Immune sensitivity |
| $y(x) = 1 - x^q$ | Immune specificity |
| $d(x) = \mu_b + \mu_i(1 - y(x))$ | Natural death rate of the host (the sum of the host's basal adult mortality rate and mortality due to autoimmunity ) |
| $\alpha(x) = \alpha_b + \alpha_i(1 - y(x))$ | Death rate due to the infection (the sum of the intrinsic virulence of the parasite and mortality due to immunopathology ) |
| $\gamma(x) = \gamma_b x$ | Recovery rate |
| $\phi(x) = \frac{\alpha(x)}{\alpha(x)+\gamma(x)+d(x)}$ | The probability of death due to the infection |
