## Supplemental Figure 1 for "The Evolution of Immune Sensitivity under Immunopathological and Autoimmune Costs"

### 2. Supplementary figures

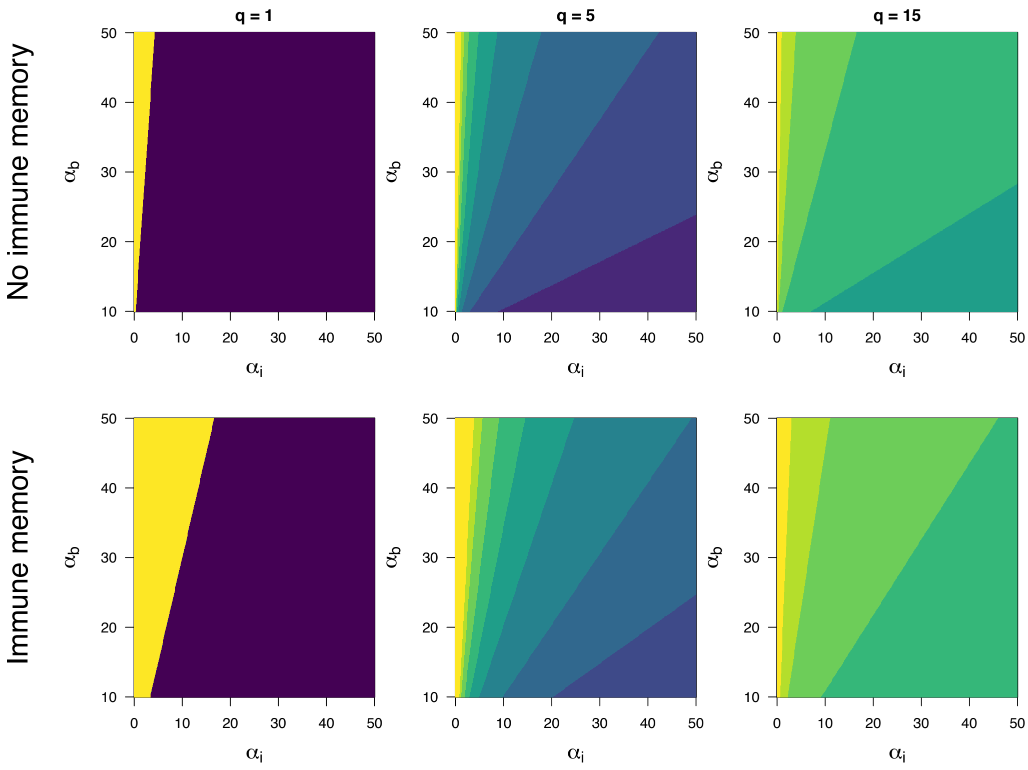


Supp. Figure 1: the effect of intrinsic parasite virulence ($\alpha_b$) and immunopathology ($\alpha_i$) on optimal levels of sensitivity, when immunity costs are immunopathological (i.e. occur during infection only). Brighter colors indicate greater optimal immune sensitivity. Results are presented for different values of parasite mimicry, with greater values of $q$ indicating a weaker trade-off between specificity and sensitivity. When $q = 1$, the tradeoff between sensitivity and specificity is linear and intermediate ESSs do not exist (selection favors either maximal sensitivity ($x = 1$) when the parasite virulence exceeds immunopathology cost (yellow regions) or maximal specificity ($x = 0$) when immunopathological costs exceed those of parasitism (purple regions)). For bigger values of $q$, the tradeoff becomes accelerating and intermediate ESS sensitivity values exist. Other parameters: $b$ = 10, $\mu_b$ = 3, $\mu_i$ = 0, $\gamma_b$ = 1.
