## Supplemental Figure 2 for "The Evolution of Immune Sensitivity under Immunopathological and Autoimmune Costs"

Supplementary materials

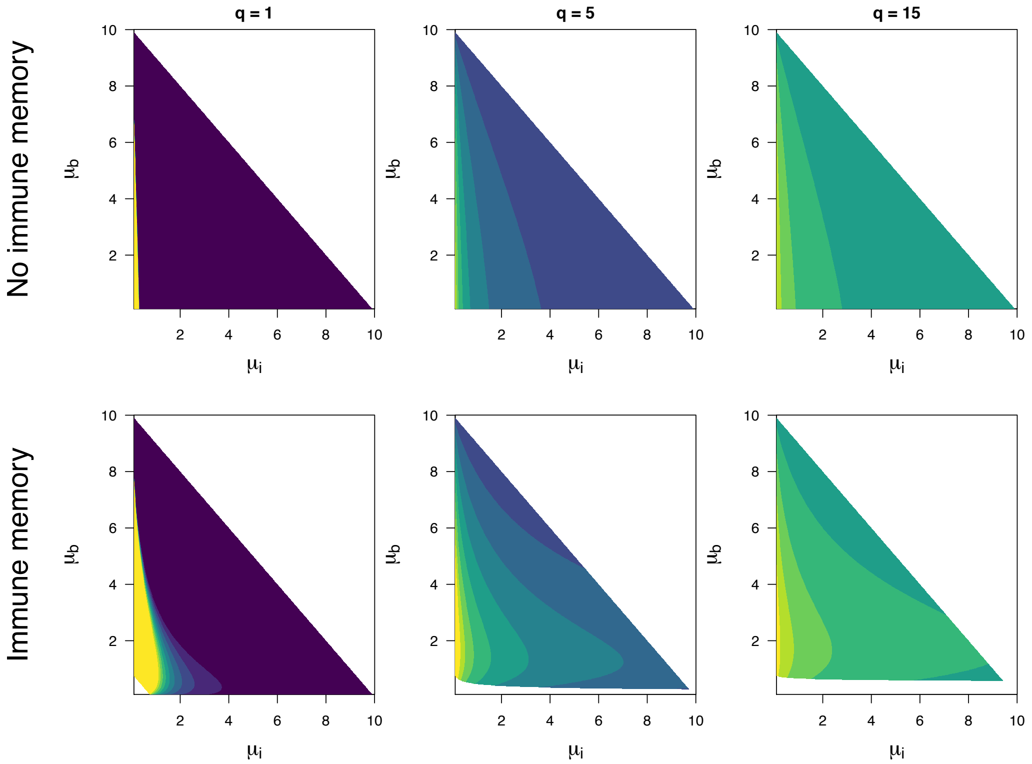


Supp. Figure 2: the effect of extrinsic host mortality ($\mu_b$) and autoimmune mortality ($\mu_i$) on optimal levels of sensitivity, when immunity costs are autoimmune (i.e. not only during infection). Brighter colors indicate greater optimal immune sensitivity. Results are presented for different values of parasite mimicry, with greater values of $q$ indicating a weaker trade-off between specificity and sensitivity. When $q = 1$, the tradeoff between sensitivity and specificity is linear and intermediate ESSs do not exist (selection favors sensitivity values close to zero (purple region) except when autoimmune mortality is very low, where sensitivity values close to one are favored (yelow region)). For bigger values of $q$, the discrimination tradeoff becomes accelerating and intermediate ESS sensitivity values exist. Other parameters: $b$ = 10, $\alpha_b$ = 3, $\alpha_i$ = 0, $\gamma_b$ = 1.
